# The wild life of ticks: using passive surveillance to determine the distribution and wildlife host range of ticks and the exotic *Haemaphysalis longicornis*, 2010-2021

**DOI:** 10.1101/2022.05.12.491673

**Authors:** Alec T. Thompson, Seth A. White, Emily E. Doub, Prisha Sharma, Kenna Frierson, Kristen Dominguez, David Shaw, Dustin Weaver, Stacey L. Vigil, Mark G. Ruder, Michael J. Yabsley

## Abstract

**Background:** We conducted a large-scale, passive regional survey of ticks associated with wildlife of the eastern U.S. Our primary goals were to better assess the current geographic distribution of exotic *H. longicornis* and to identify potential wild mammalian and avian host species. However, this large-scale survey also provided valuable information regarding the distribution and host associations for many other important tick species that utilize wildlife as hosts.

**Methods:** Ticks were opportunistically collected by cooperating state and federal wildlife agencies. All ticks were placed in the supplied vials and host information was recorded, including host species, age, sex, examination date, location (at least county and state), and estimated tick burden were recorded. All ticks were identified to species using morphology, suspect *H. longicornis* were confirmed through molecular techniques.

**Results:** In total, 1,940 hosts were examined from across 369 counties from 23 states in the eastern U.S. From these submissions, 20,626 ticks were collected and identified belonging to 11 different species. Our passive surveillance efforts detected exotic *H. longicornis* from nine host species from eight states. Notably, some of the earliest detections of *H. longicornis* in the U.S. were collected from wildlife through this passive surveillance network. In addition, numerous new county reports were generated for *Amblyomma americanum, A. maculatum, Dermacentor albipictus, D. variabilis*, and *Ixodes scapularis*.

**Conclusions:** This study provided data on ticks collected from animals from 23 different states in the eastern U.S. between 2010 – 2021 with the primary goal of better characterizing the distribution and host-associations of the exotic tick *H. longicornis*; however new distribution data on tick species of veterinary or medical importance was also obtained. Collectively, our passive surveillance has detected numerous new county reports for *H. longicornis* as well as *I. scapularis*. Our study utilizing passive wildlife surveillance for ticks across the eastern U.S. is an effective method for surveying a diversity of wildlife host species allowing us to better collect widespread data on current tick distributions relevant to human and animal health.

## Background

Ticks and tick-borne diseases constitute a major threat to human and animal health and are rapidly becoming recognized as a global One Health issue. Numerous underlying factors such as climate change, habitat fragmentation, and increases in globalization with the movement of humans and animals to new areas of the world all promote the geographic expansion of multiple tick species and their pathogens [1–4]. The spread of non-native parasites is a significant concern for disease emergence and native species conservation; therefore it is of extreme importance to identify these exotic ticks and their pathogens and take effective steps to prevent their dispersal and establishment, of which presents enormous challenges to both conservation and international commerce [5–7]. In the case of exotic ticks, the detection and management of these species often fail for a variety of reasons resulting from their unique and often complex life history traits and ability to utilize a variety of domestic, livestock, and wildlife hosts.

Passive surveillance is a commonly used method by health officials and researchers to investigate the geographic distribution and host associations of ticks. Many of these passive surveillance strategies involve image submissions of ticks for expert, artificial intelligence, or crowd sourced identification [8–11], the use of electronic patient records from companion animals [12], and most commonly whole tick submissions from citizen scientists, veterinarians, and physicians [13–18]. Only a few published studies using passive surveillance have included ticks collected from wildlife hosts, and of those studies, most were to statewide surveys leaving gaps in our understanding of the regional distribution of ticks relevant to both human and animal health [19–23].

A tick of recent One Health significance in the United States is the Asian longhorned tick, *Haemaphysalis longicornis* (Acari: Ixodidae). Native to East Asia, *H. longicornis* has become invasive to several regions of the world including Australia, New Zealand, and most recently the U.S., having first been detected outside of quarantine zones in 2017 on an Icelandic ewe (*Ovis aries*) from New Jersey [24–26]. Subsequent reexaminations of archived specimens revealed the presence of *H. longicornis* in the U.S. on several wildlife species from multiple states dating as early as 2010 on a white-tailed deer (*Odocoileus virginianus*) from West Virginia (present study) [27]. This case of *H. longicornis* on wildlife almost a decade prior to the tick being detected on livestock justifies a dire need for a comprehensive tick survey of wildlife hosts within the U.S.

Within the established range, *H. longicornis* infests a variety of mammalian and avian species (including companion animals, livestock, and wildlife) and is found in a variety of geographic and climatic habitats [25–28]. Since the initial discovery outside of quarantine zones, *H. longicornis* has now been detected in 17 states and has become an increasing human and veterinary health concern as it is either suspected or confirmed to be a vector for several pathogens. Recent laboratory infection trials have indicated *H. longicornis* as a competent vector for *Rickettsia rickettsii*, the causative agent for Rocky Mountain Spotted Fever, and Heartland virus, but experimentally was not a suitable vector for *Borrelia burgdorferi* sensu strico or *Anaplasma phagocytophilum* Ap-Ha, the causative agents for Lyme Disease and Human Granulocytic Anaplasmosis, respectively [29–32]. However, despite the experimental transmission studies, several medically important pathogens have been detected in environmentally collected host-seeking *H. longicornis* including *B. burgdorferi, A. phagocytophilum* Ap-Ha (both detected in populations from Pennsylvania [33,34]), and *Rickettsia felis* and Bourbon virus (detected in populations from Virginia [35,36]). Of veterinary importance, native genotypes of the white-tailed deer variant of *Anaplasma phagocytophilum* (Ap-1) and a *Hepatozoon* sp. have been detected in host-seeking *H. longicornis* from Virginia [35]. Additionally, *H. longicornis* is a confirmed vector for an exotic protozoan parasite, *Theileria orientalis* Ikeda, the cause of cattle mortalities at a farm in Virginia although infections have been noted to be more widespread in Virginia and West Virginia [37–39]. Finally, there have been multiple reports of intense infestations on cattle resulting in exsanguination events in North Carolina and previous studies report severe *H. longicornis* infestations on wildlife species [40–42].

Currently published surveillance studies for *H. longicornis* in the U.S. are limited geographically and as a result are unlikely to capture the potential wildlife host range utilized by *H. longicornis* [35,40,41,43–45]. In addition, habitat suitability models primarily focusing on climatic and geographic variables to predict the potential range of *H. longicornis* have been reported, but they were built around limited datasets of *H. longicornis* occurrences (rather than established population data) and therefore may not accurately depict all suitable habitats in the U.S. [46–48]. In this study, we conducted a large-scale, passive regional survey of ticks associated with wildlife of the eastern U.S. Our primary goals were to better assess the current geographic distribution of exotic *H. longicornis* and to identify potential wild mammalian and avian host species. However, this large-scale survey also provided valuable information regarding the distribution and host associations for many other tick species of medical and veterinary importance that utilize wildlife as hosts.

## Materials and Methods

Wildlife host surveillance for *H. longicornis* started in fall 2017 after the initial detection of this tick outside of quarantine zones in New Jersey and is currently ongoing [24,49]. Tick collections kits consisting of 15mL vials pre-filled with 70% ethanol, forceps, collection instructions, blank labels, and datasheets were shared with state and federal wildlife agencies that were members of the Southeastern Cooperative Wildlife Disease Study (SCWDS) and with states currently reporting *H. longicornis* infestations. Participating agencies were asked to disseminate tick collection kits and instructions to agency staff members. Ticks were collected from a variety of sources including wildlife during health surveys, car strike kills, nuisance animal removal, hunter checks, or during sample collection for other ongoing studies. All ticks were placed in the supplied vials and general information such as host species, age, sex, examination date, location (at least county and state), and estimated tick burden was recorded. Ticks and corresponding data sheets were then submitted to SCWDS for identification. In addition, ticks were also opportunistically collected from diagnostic case submissions to SCWDS from member states.

Upon receipt, tick vials were given a unique identification number and screened for *Haemaphysalis* spp. ticks using morphology [50]. All suspect *H. longicornis* were examined by at least two people and were confirmed using polymerase chain reaction (PCR) targeting the 16S rRNA gene, analyzed using restriction fragment length polymorphisms (RFLPs), and sequenced as described by Thompson et al. [51]. Specimens of *H. longicornis* collected from either a new host, new county, or new state were submitted to the National Veterinary Sciences Laboratory (NVSL) in Ames, Iowa for morphologic confirmation and archiving purposes. All other ticks in the vials were identified to species using morphology keys [50,52–56]. Specimens that were damaged or in poor condition (i.e., missing capitulum or eviscerated) were identified to genus and life stage when possible. The data collected from this surveillance effort were reported to USDA-APHIS and presented in the monthly National *Haemaphysalis longicornis* (Asian longhorned tick) Situation Report [49]. A subset of these data were included in summary form in a previous report on the distribution of *H. longicornis* [27]. Established tick populations for county level data were classified based on the Centers for Disease Control and Prevention (CDC) criteria of ≥6 individual ticks or >1 life stage collected in a span on one year, where ticks collected from deer or small- or medium-sized mammals being acceptable to classify a county status [57,58]. Classification of any new established counties was compared with previous data when available (National Arboviral Surveillance System ArboNET; https://www.n.cdc.gov/Arbonet/) [59].

Since 1961, SCWDS has assisted various state and federal agencies conduct herd health checks on white-tailed deer. As part of this work, ticks were collected and identified. These archived SCWDS tick specimens from 2010 – 2017 were also included in the study. No ticks included in this study were tested for pathogens because most host-attached ticks could contain host blood convoluting pathogen detection data (i.e., was the tick infected or was the host infected).

## Results

In total, 1,940 hosts were examined from across 369 counties from 23 states in the eastern U.S. (Supplemental Table 1, Figure 1). Although agencies were asked to collect from wildlife, a few submissions from domestic animals and people were submitted so these were also included in our study. White-tailed deer was the most sampled host species (n = 1,371; 71%) followed by black bear (*Ursus americanus*; n = 226; 12%), elk (*Cervus canadensis*; n = 96; 5%), mule deer (*Odocoileus hemionus*; n = 66; 3%), and wild turkey (*Meleagris gallopavo*; n = 42; 2%). Other wildlife species making up the remaining ten percent of hosts sampled included: woodchuck (*Marmota monax*; n = 22), spotted skunk (*Spilogale putorius*; n = 21), raccoon (*Procyon lotor*; n = 16), coyote (*Canis latrans*, n = 11), domestic dog (*Canis familiaris*; n = 11), wild pig (*Sus scrofa*; n = 11), bobcat (*Lynx rufus*; n = 9), red fox (*Vulpes vulpes*; n = 6), Virginia opossum (*Didelphis virginiana*; n = 6), striped skunk (*Mephitis mephitis*; n = 5), gray fox (*Urocyon cinereoargenteus*; n = 2), and white-footed mouse (*Peromyscus leucopus*; n = 2). We received one submission each from a desert bighorn sheep (*Ovis canadensis nelson*), Northern bobwhite (*Collinus virginianus*), red-backed vole (*Clethrionomys* sp.), red deer, (*Cervus elaphus*), ruffed grouse (*Bonasa umbellus*), and woodland jumping mouse (*Napaeozapus insignis*). Ticks were received from single cattle herd (*Bos taurus*); however individual infested cattle were not counted. Finally, two submissions were received from humans and eight submissions were collected from an environmental source (Supplemental Table 1).

**Figure 1.**
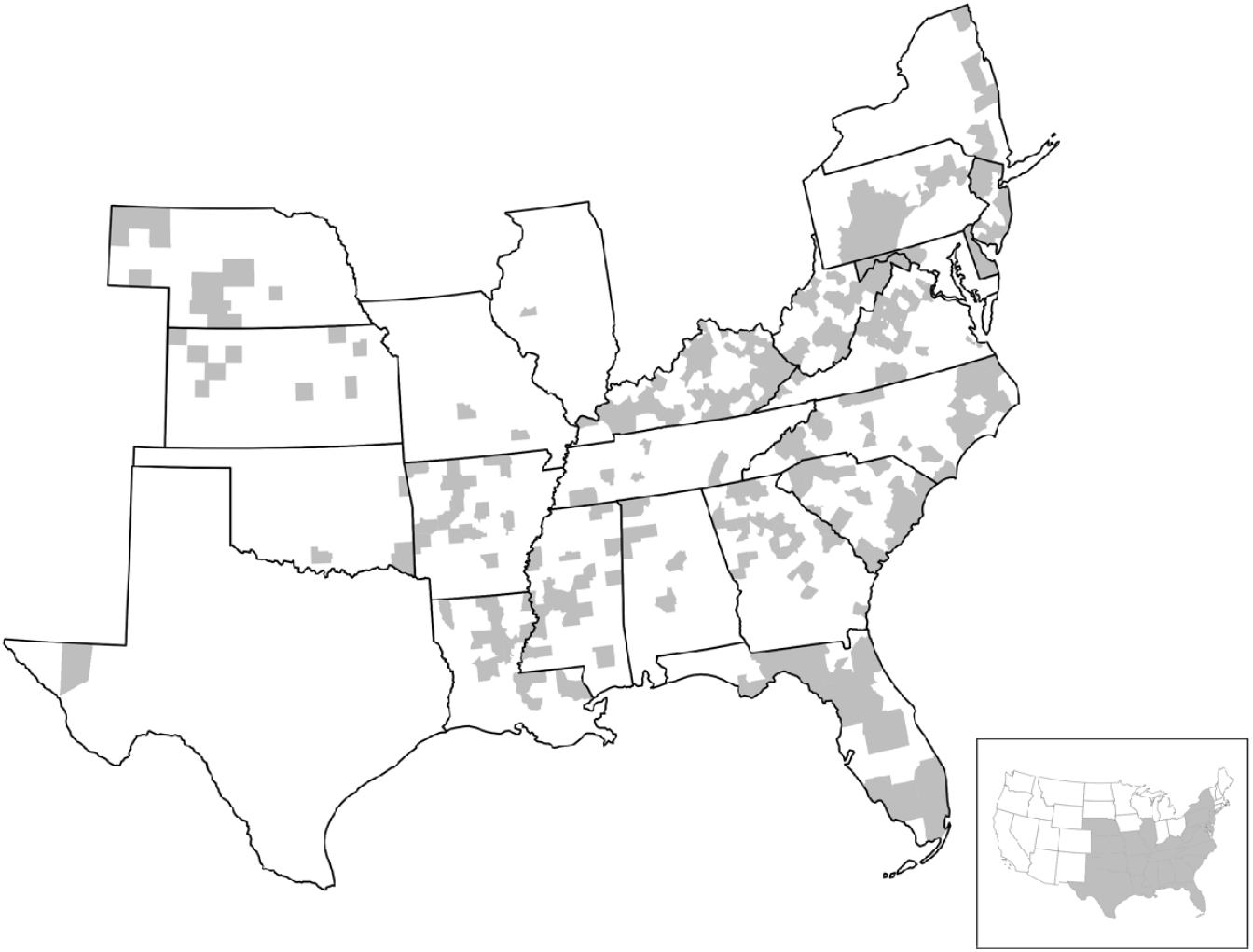
Agency submissions, 2010-2021. Counties highlighted in grey indicate at least one tick submission was received during the study. Inset represents states that submitted samples.

From these submissions, 20,626 ticks were collected and identified. *Amblyomma americanum* (n = 10,942; 53%) and *Ixodes scapularis* (n = 4,846; 24%) were the two most collected species followed by *Dermacentor variabilis* (n = 1,804; 9%), *Dermacentor albipictus* (n = 1,240; 6%), *Amblyomma maculatum* (n = 848; 4%), and *H. longicornis* (n = 451; 2%). The remaining two percent of tick species collected included: *Ixodes cookei* (n = 201), *Ixodes affinis* (n = 38), *Otobius megnini* (n = 26), *Ixodes texanus* (n = 15), and *Haemaphysalis leporispalustris* (n = 1). A small number of ticks (one *Amblyomma* and 202 *Ixodes*) could not be identified to species due to missing morphologic features (Supplemental Table 1).

Our passive surveillance efforts detected the exotic *H. longicornis* from 9 host species from 8 states including Georgia, Kentucky, Maryland, New Jersey, North Carolina, Pennsylvania, Virginia, and West Virginia (Supplemental Table 1, Figure 2F). White-tailed deer (n = 41) and elk (n = 12) were the two host species most frequently detected with *H. longicornis* infestations followed by domestic dog (n = 4), black bear (n = 2), and coyote (n = 2), with human, red fox, and Virginia opossum each having one detection (Figure 3). Individual cows were not counted in this study; however, a herd in Pickens County, GA was found to be infested with *H. longicornis*. Two submissions from environmental sources were also positive for *H. longicornis*. The passive surveillance resulted in many of the first county, state, and host reports for *H. longicornis* in the U.S. (Table 1). Notably, some of the most historical detections of *H. longicornis* in the U.S. were collected from wildlife through the SCWDS passive surveillance network. The earliest being a white-tailed deer from West Virginia in 2010 (previously misidentified as *H. leporispalustris*) as well as a black bear from Kentucky, a white-tailed deer from West Virginia, and a Virginia opossum from North Carolina in 2017 (additional specimens previously misidentified as *H. leporispalustris*). Based on CDC guidelines for established tick populations (≥6 ticks or >1 life stage collected in a span of one year), 11 counties were classified as established, nine of which represent new counties including the furthest south detection of *H. longicornis* in the U.S. from Pickens County, GA [59] (Figure 2F)

**Figure 2.**
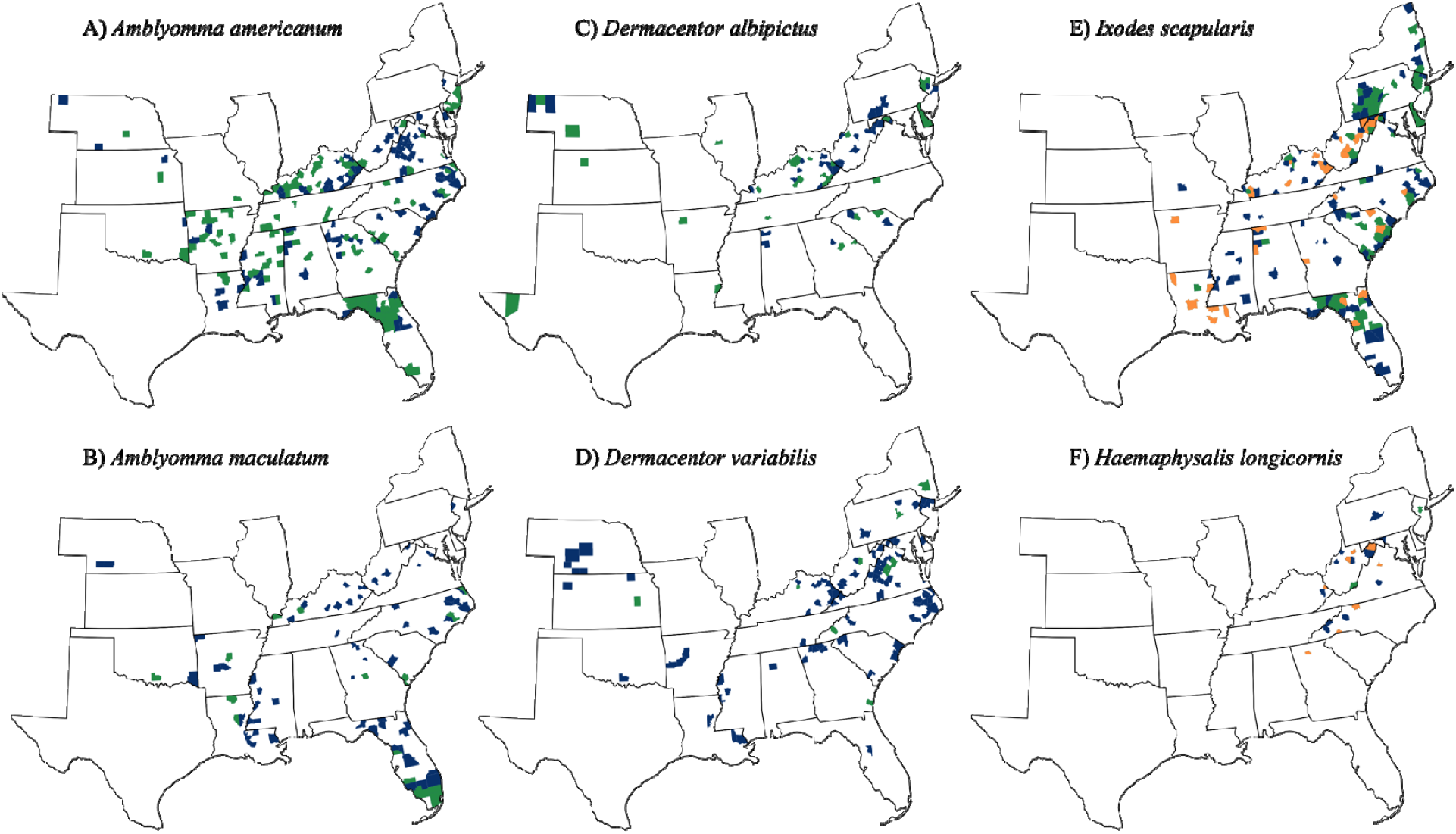
Spatial data for individual tick species detected from surveillance between 2010 – 2021. Green counties indicate an established population as defined by the CDC. Orange counties indicate a new established classification for a county. Blue counties represent other detections made from the present study. A) *Amblyomma americanum*; B) *Amblyomma maculatum*; C) *Dermacentor albipictus*; D) *Dermacentor variabilis*; E) *Ixodes scapularis*; F) *Haemaphysalis longicornis*.

**Figure 3.**
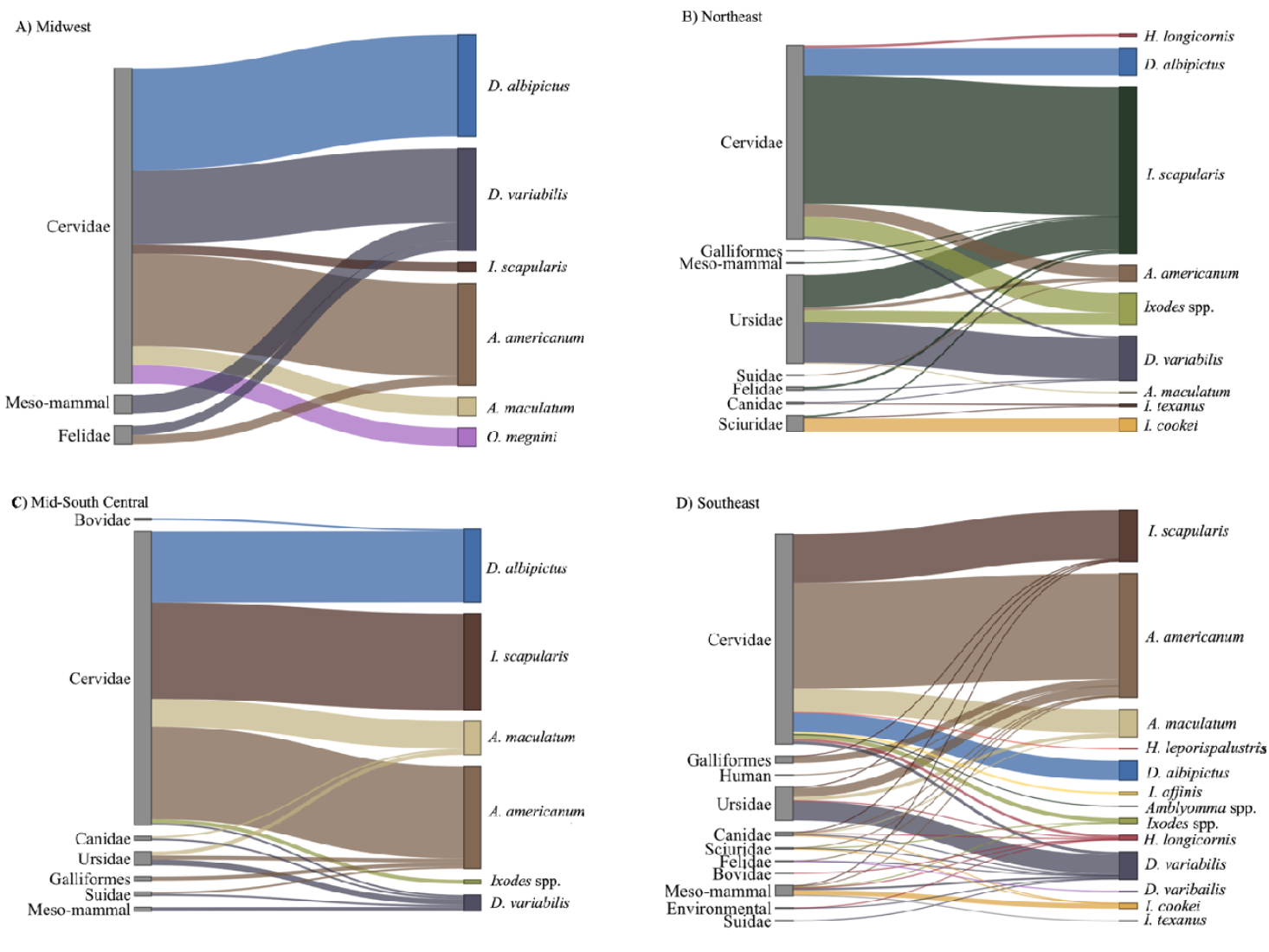
Tick-host associations by region. For all panels, left axis represents hosts and right axis represents tick species, links between the two axes depicts the tick species that was collected from the different host groups. A) Midwest (includes IL, IN, KS, MO, NE, OH); B) Northeast (includes CT, DE, MA, MD, NJ, NY, PA, RI); C) Mid-South Central (includes AR, LA, OK, TX); D) Southeast (includes AL, FL, GA, KY, MS, NC, SC, TN, VA, WV).

**Table 1.** First host, county, or state detections of *Haemaphysalis longicornis* through this passive wildlife agency surveillance effort, 2010 – 2021

| Date | State | County | Host<sup>a</sup> | Detection<sup>b</sup> | NVSL<sup>c</sup> Accession |
| --- | --- | --- | --- | --- | --- |
| 8/13/10 | West Virginia | Tyler | White-tailed deer | S, C, H | 11-015712 |
| 6/2/17 | Kentucky | Floyd | Black bear | S, C, H | 20-011466 |
| 9/1/17 | West Virginia | Taylor | White-tailed deer | S, C | 18-021706 |
| 9/15/17 | North Carolina | Polk | Virginia opossum | S, C, H | 17-032017 |
| 3/28/18 | North Carolina | Madison | Domestic dog | C | 19-036810 |
| 5/27/18 | New Jersey | Somerset | White-tailed deer | C |  |
| 6/22/18 | West Virginia | Upshur | Coyote | C, H |  |
| 5/25/18 | Maryland | Washington | White-tailed deer | S, C | 18-025265 |
| 7/10/18 | Pennsylvania | Centre | White-tailed deer | S, C | 18-025264 |
| 7/11/18 | Virginia | Greene | Red fox | C, H | 19-005067 |
| 7/15/18 | Kentucky | Martin | Elk | C, H | 19-006599 |
| 8/1/18 | West Virginia | Hampshire | White-tailed deer | C | 19-012661 |
| 8/28/18 | West Virginia | Wirt | White-tailed deer | C | 19-012659 |
| 8/30/18 | West Virginia | Lewis | White-tailed deer | C |  |
| 10/3/18 | West Virginia | Grant | White-tailed deer | C |  |
| 4/22/19 | North Carolina | Haywood | Domestic dog | C | 19-036811 |
| 7/18/19 | North Carolina | Surry | White-tailed deer | C |  |
| 8/27/19 | West Virginia | Mineral | White-tailed deer | C | 20-006056 |
| 9/2/20 | North Carolina | Caldwell | White-tailed deer | C | 20-035011 |
|  | West Virginia | Monroe | White-tailed deer | C |  |
| 7/21/21 | Virginia | Appomattox | Black bear | C | 21-035736 |
| 9/21/21 | Georgia | Pickens | Cow | S, C |  |

<sup>a</sup> Black bear (*Ursus americanus*), domestic dog (*Canis familiaris*), elk (*Cervus canadensis*), red fox (*Vulpes vulpes*), white-tailed deer (*Odocoileus virginianus*)
<sup>b</sup> Indicates first or earliest detection for state (S), county (C), or host (H)
<sup>c</sup> NVSL, National Veterinary Services Laboratory

Of the five most abundant tick species collected, the most common species, *A. americanum*, was detected from 18 of the 23 sampled states and was found to be established in 115 different counties. It was not reported from the most northeastern, midwestern and southwestern regions of the sampled area (Figure 2A). Eleven different mammalian species and one avian species were infested with *A. americanum. Ixodes scapularis* was the second most common tick species collected and had a broad range. This tick was detected from all states that submitted specimens except for the most western sampled states (Nebraska, Kansas, Oklahoma, and Texas) and Illinois and classified as established in 91 different counties which included 34 new county reports (ArboNET). *Ixodes scapularis* was collected from 14 mammalian host species and two avian host species. Similar to *I. scapularis*, both *D. albipictus* and *D. variabilis* had broad distributions however they were detected less frequently and from four mammalian host species and 14 mammalian host species, respectively. *Amblyomma maculatum* was detected primarily in the more southern and coastal states sampled on six mammalian host species, however we also received specimens from counties outside of its previously reported range in central Kentucky, Nebraska, New Jersey, Virginia, and West Virginia [60]. (Figure 2, Figure 3). *Ixodes affinis* was identified from a small number of white-tailed deer from Florida and North Carolina. The spinose ear tick, *O. megnini*, was rarely detected in submissions from the Midwest. Finally, *I. cookei* and *I. texanus* were periodically collected when submitted ticks from medium-sized mammals (Supplemental Table 1, Supplemental Table 2).

## Discussion

This study provided data on ticks collected from animals from 23 different states in the eastern U.S. between 2010 – 2021 with the primary goal of better characterizing the distribution and host-associations of the exotic tick *H. longicornis*. However, new data for several native tick species of veterinary or medical importance were also obtained. Collectively, our passive surveillance has detected numerous new established county reports for *H. longicornis* as well as *I. scapularis*. Over 1900 wildlife and domestic hosts were sampled representing 23 mammalian and three avian species, however a majority of these hosts were from the families Cervidae (elk, mule deer, red deer, white-tailed deer; n = 1,534) and Ursidae (black bear; n = 226), as wildlife agency personnel are often in close contact with these species. However, it is important to note, this sampling bias skews the observed diversity and collection frequency to overrepresent those tick species and life stages that are most likely to feed on cervids and bears. In addition, 206 medium sized mammals (bobcat, coyote, domestic dog, gray and red fox, raccoon, spotted and striped skunk, Virginia opossum, and woodchuck) were also sampled allowing some insight into the tick-host associations of these species. Our study utilizing passive wildlife surveillance for ticks across the eastern U.S. is an effective method for surveying a diversity of wildlife host species allowing us to better collect widespread data on current tick distributions relevant to human and animal health.

Wildlife are important hosts for many tick species as they can serve as maintenance hosts and potential disseminators, and, in some cases, wildlife species have facilitated increases in the range of tick species either through natural movements, migration, or human-facilitated translocation [2,4,61]. For example, the increasing populations of white-tailed deer and wild turkey in the eastern U.S. have been linked to the increasing abundance and broader distribution of *A. americanum* [62], migratory birds have been implicated in the expansion of the range of *I. scapularis* [61], and the movement of large carnivores and domestic animals have been associated with the gradual northern expansion of *D. variabilis* [63]. Finally, both migratory birds and medium to large mammal species have been suggested to facilitate the expansion of *A. maculatum* [64,65]. For exotic *H. longicornis*, it is still debated how this tick is spreading within the U.S., though its initial introduction to the U.S. is believed to be caused via domestic animal and livestock movement [66,67]. Furthermore, several of the tick species detected in this study are more common on wildlife than domestic species and are rarely detected via environmental detections [68]. For example, given certain life history traits *D. albipictus, H. leporispalustris, I. cookei, I. texanus*, and *O. megnini* would likely not be detected in surveillance studies focusing on domestic animals or utilizing only environmental sampling. In addition, many wildlife species serve as reservoir hosts for many tick-borne pathogens relevant to human and animal health. By characterizing tick distributions via wildlife host sampling, we can begin to better predict areas of higher disease risk where vector and reservoir host co-occur.

Per current CDC guidelines, for a tick population to be classified as “established”, it requires the collection of ≥6 ticks or >1 life stage within a single year, either from the environment or from deer and small- to medium-sized mammals, all else is considered “reported” [57,69]. For our county classifications, we considered deer to include any cervid species and small- to medium-sized mammals to include anything smaller than a coyote. In general, the distributions of ticks and their host-associations detected in this study were similar to what has been previously reported in the literature [10,18,19,57,64,70–74]. Unfortunately, for certain tick species (e.g., *A. americanum, A. maculatum, D. albipictus*, and *D. variabilis*) there are currently limited data designating counties as reported vs. established counties, therefore we were unable generate any new established county data for these species. However, with significantly more vector-borne disease cases being reported in the U.S. and emphasis on surveillacnce, we expect similar large-scale datasets like the ArboNET *I. scapularis* data to become available [75].

Two species of *Haemaphysalis* were detected: the exotic Asian longhorned tick *H. longicornis* (n=451), and the native rabbit tick, *H. leporispalustris* (n=1). Several studies that have been conducted in the U.S. after the initial detection of *H. longicornis* have documented a broad host range for *H. longicornis*; however, these studies were each focused in relatively small geographic areas (e.g., Connecticut, Pennsylvania, New Jersey, New York, or Virginia) [33,35,43,45,59]. Our passive geographically broad-scale surveillance for *H. longicornis* was an effective method for providing new and rapid data on the distribution of this tick in the U.S. The passive surveillance efforts detected *H. longicornis* from nine host species (black bear, cow, coyote, domestic dog, elk, human, red fox, Virginia opossum, and white-tailed deer) from eight different states and resulted in some of the first county, state, and host collections in the U.S. Although these data are compiled in the current study for a long study period, data were reported to the USDA in real-time for inclusion in monthly situation reports [49]. When this tick was first detected in 2017, several studies were conducted to determine if wildlife were infested and retrospective data were either reviewed or archived ticks were examined, some of which were included in this study. The single *H. leporispalustris* was collected from a white-tailed deer in Georgia. Although, this tick is commonly associated with lagomorphs and avian species, detections of this tick on deer is not uncommon; however, this highlights the importance of being able to distinguish these two morphologically similar *Haemaphysalis* spp. [74].

This study documents the earliest record of *H. longicornis* in the U.S. which was collected from a white-tailed deer in 2010 in West Virginia, years before it was identified as established in North America [27]. Additionally, other SCWDS-led tick surveillance efforts conducted during 2017, the same year it was intuially detected in New Jersey, collected *H. longicornis* from additional wildlife species and new states (black bear from Kentucky and Virginia opossum from North Carolina), demonstrating the important role of wildlife surveillance in detecting ticks. Combined, this data indicates that *H. longicornis* was present in the U.S. for years before its detection in New Jersey and was much more widespread that initially believed. Initially, *H. longicornis* was primarily detected in the mid-Atlantic states, where this tick is now well known to occur, however, our passive surveillance efforts detected *H. longicornis* as far north as Pennsylvania and as far south as Georgia and have contributed greatly to the current understanding of this tick’s current geographic and host ranges [49]. In total from these data, 11 counties were classified as established for *H. longicornis*, nine of which represent new counties including the one for the most southern detections of *H. longicornis* in the U.S.—Pickens County, GA [59].

During this study, four *Ixodes* species were collected from wildlife. As expected, given the hosts sampled, *I.scapularis* (n = 4,846) was the most abundant and widespread *Ixodes* species detected. This tick species was not detected in the more western states, but this is likely due to fewer submitted samples from that area and these states being on the edge of currently recognized *I. scapularis* distribution [57]. From our surveillance, a total of 35 new counties have established *I. scapularis* populations (ArboNET). These new counties had either been previously classified as reported or had no data. Our passive surveillance using wildlife has provided valuable information for public health officials as *I. scapularis* is a vector for numerous important human pathogens such as *Borrelia burgdorferi* and *Babesia microti*. Surprisingly, few *I. affinis* ticks (n = 38) were detected during this study with positive hosts being white-tailed deer submitted from Florida and North Carolina. *Ixodes affinis* is widespread in the coastal regions of the southeastern U.S. and currently undergoing a northern range expansion [64]. Unlike, *I. scapularis, I. affinis* is found on deer during the summer so the bias of deer sampling during hunting season likely explains the limited detections. Additionally, *I. cookei* (n = 201) and *I. texanus* (n = 15) were also collected and both are widespread east of the Mississippi River and mainly infest a diversity of small- to medium-sized hosts. Our detection of *I. cookei* on a red-backed vole represents a new host record for this tick [74]. Finally, 202 *Ixodes* ticks could not be identified due to damage to key morphological features, these samples represent a limitation to passive surveillance work as the quality of specimens submitted may not always be ideal. Fortunately, molecular techniques are available to identify these specimens; however. because the overall number of damaged ticks received was low (<1%), molecular identification was not pursued.

Two species of *Amblyomma* ticks were collected during this study: *A. americanum* (n = 10,942) and *A. maculatum* (n = 848). *Amblyomma americanum* was the most abundant tick species collected in the study, however both *A. americanum* and *A. maculatum* were widespread in the southern states with detections becoming more limited further north toward the edge of their currently recognized ranges [64,73]. Surprisingly, we detected *A. maculatum* more inland than previously reported, however, with a lack of publicly available county-level data, we are unable to determine if our surveillance for these species has resulted in any new distribution records. Regardless, 115 and 16 counties were classified as established for *A. americanum* and *A. maculatum*, respectively. It is important to note, that these two species are important pathogen vectors in the southeastern U.S. and are more likely to transmit pathogens to domestic animals and humans than *I. scapularis* due to their aggressive host-biting behaviors [62,64]. In addition, *A. americanum* is most commonly associated with alpha-gal syndrome (red meat allergy) [76].

Finally, two *Dermacentor* ticks were also collected during this study, *D. variabilis* (n = 1,804) was the most abundant followed by *D. albipictus* (n = 1,240). Both species of tick were sporadically detected but collected widely across the study region. Submitted samples resulted in 11 and 35 established counties for *D. variabilis* and *D. albipictus*, respectively. *Dermacentor variabilis* is a vector for *Rickettsia rickettsii*, the causative agent of Rocky Mountain Spotted Fever, and commonly found infesting medium- to large-sized mammals as adults and smaller mammals during its immature life stages [74]. Black bear was the host most commonly infested with this tick species. Additionally, *D. albipictus*, known as the winter tick, is commonly associated with cervid species, which is consistent with our detections. *Dermacentor albipictus* is a serious pest for moose, on which severe infestations can lead to alopecia, emaciation, and potentially death. Interestingly, infestations rarely become as severe on other cervid species; however, recently an elk in Pennsylvania was found dead due to severe *D. albipictus* infestations [77].

A majority of ticks collected from this study were hard ticks (Ixodidae), but we also detected 26 soft ticks, *Otobius megnini* (Argasidae). Soft ticks are rarely found in the environment as they are mostly nidicolous (nest-dwelling) [78]. *Otobious megnini* is widely distributed in the southern and western U.S., however sporadic detections have been made in the eastern U. S. as a result of animal movement [74,79,80]. The limited detections in our study were likely because this tick lives in the ear canal of its hosts and biologist collaborators were only requested to examine the skin of animals for ticks. This tick is currently not known to transmit any pathogens, but it can infest a diversity of wildlife and domestic mammalian hosts [74].

This study provided important new distribution and host data for many tick species, but there are several limitations regarding the utility and interpretation of the data as highlighted in Eisen and Eisen [81]. Our tick collections were opportunistic and based on the ability of our agency partners to collect ticks from the hosts; therefore, we accurate data on tick burdens on each host. In addition, since a majority the ticks collected in this study were collected from hosts, it is challenging to interpret the precise spatial data (when and if provided), especially for hosts that have large home ranges as we do not know the origin of the tick or the precise location where it interacted with the hosts. Another limitation is that no ticks included in this study were tested for pathogens because most host-attached ticks could contain host blood which can convolute pathogen detection data (i.e., was the tick infected or was the host infected). To effectively determine pathogen prevalence and distribution, testing of host-seeking ticks is recommended [58,81]. Even with these drawbacks, this type of large-scale study and the data it generated provided valuable baseline data for many new hosts and regions for ticks of medical and veterinary concern.

## Conclusions

Our study utilizing passive wildlife surveillance for ticks across the eastern U.S. is an effective method for surveying a diversity of wildlife host species allowing us to better collect widespread data on current tick distributions relevant to human and animal health. There are currently a few recent large scale tick surveillance studies in the U.S., however none focus primarily on wildlife species. This study has collected valuable data regarding the distribution and host associations of exotic *H. longicornis*, in addition, valuable information about native tick species relevant to human and animal health was also collected.

## Declarations

### Ethics approval and consent to participate

Not applicable

### Consent for Publication

All authors consent to the publication of this research study.

### Availability of data and materials

The datasets used and analyzed during the current study are available from the corresponding author on reasonable request.

### Authors’ contributions

AT, ED, DS, MR, MY, SV, and SW participated in study design and contacted agency partners. AT and MY drafted manuscript. AT analyzed and visualized all data. All authors collected data and read and approved the final manuscript.

## Acknowledgements

We thank numerous biologists and veterinarians in SCWDS member state agencies for sample collection and submission.

---

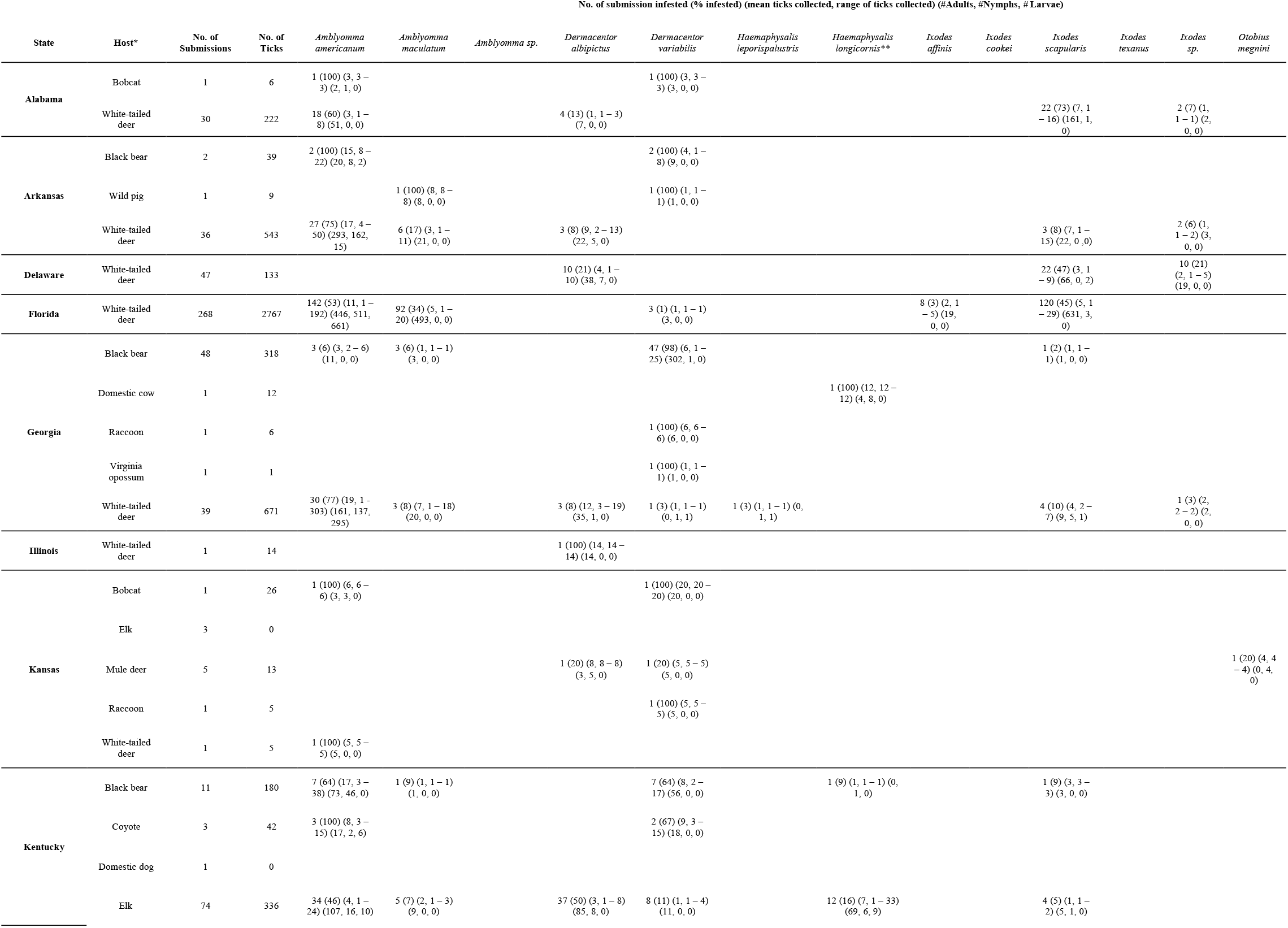
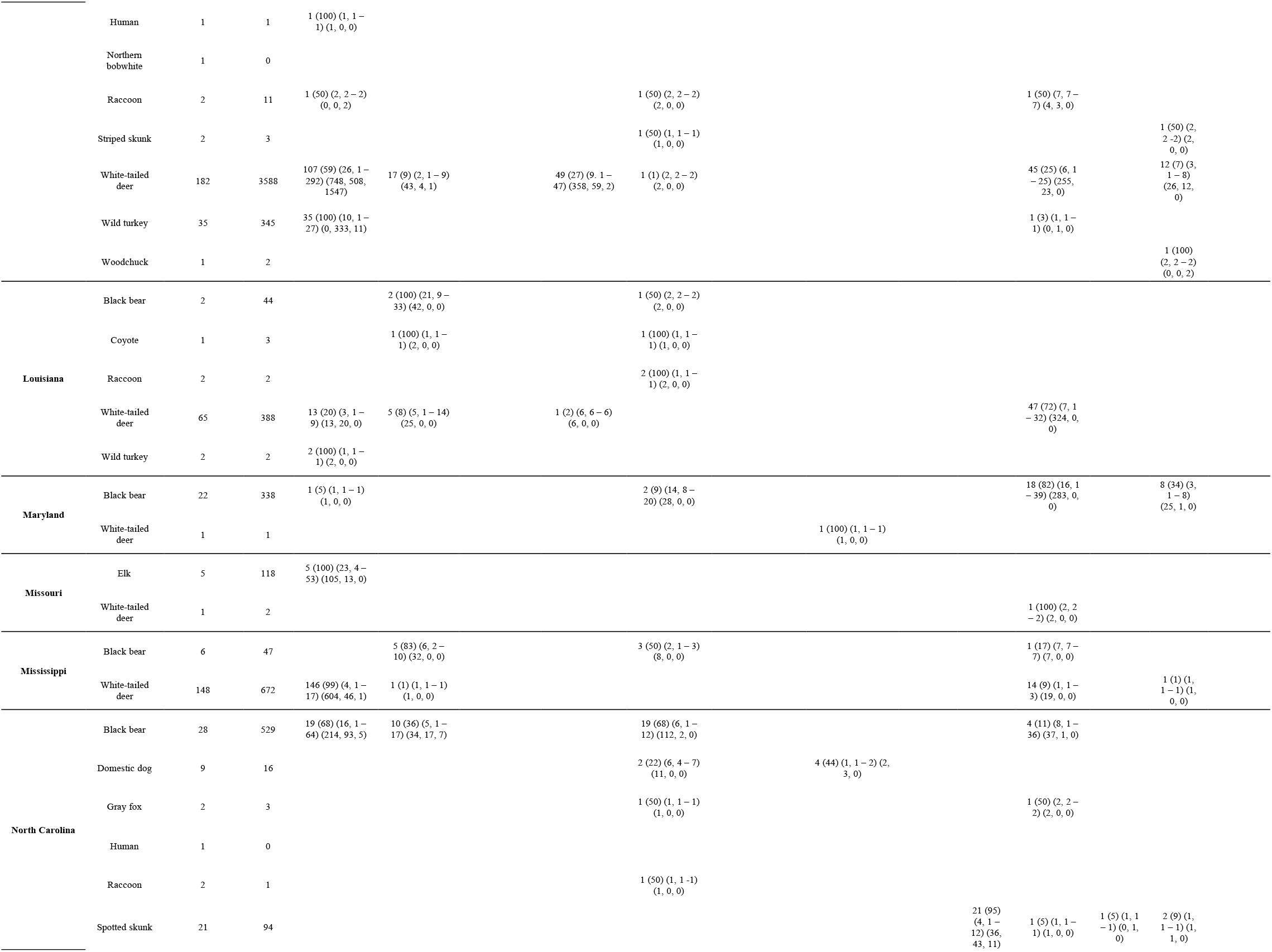
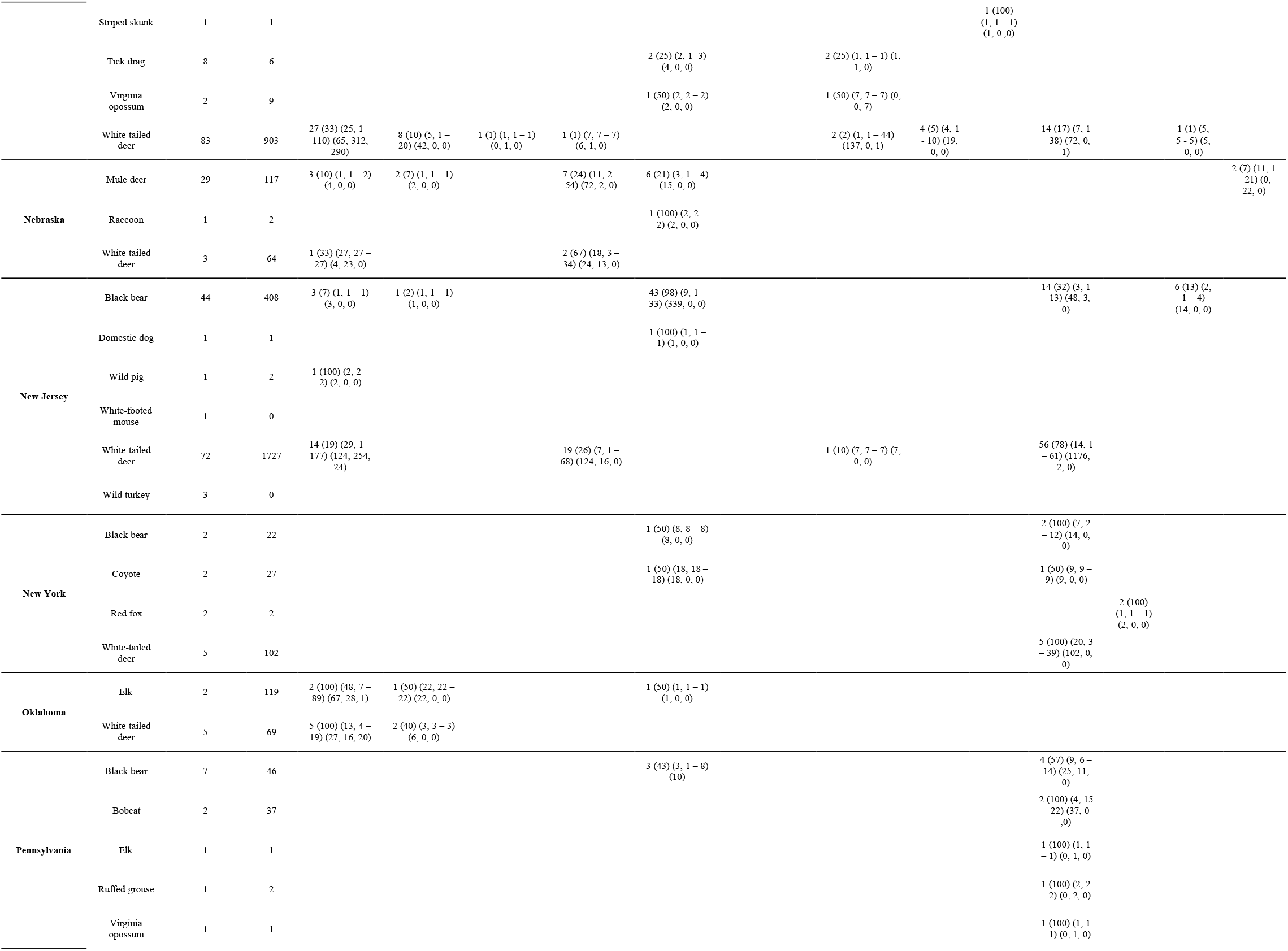
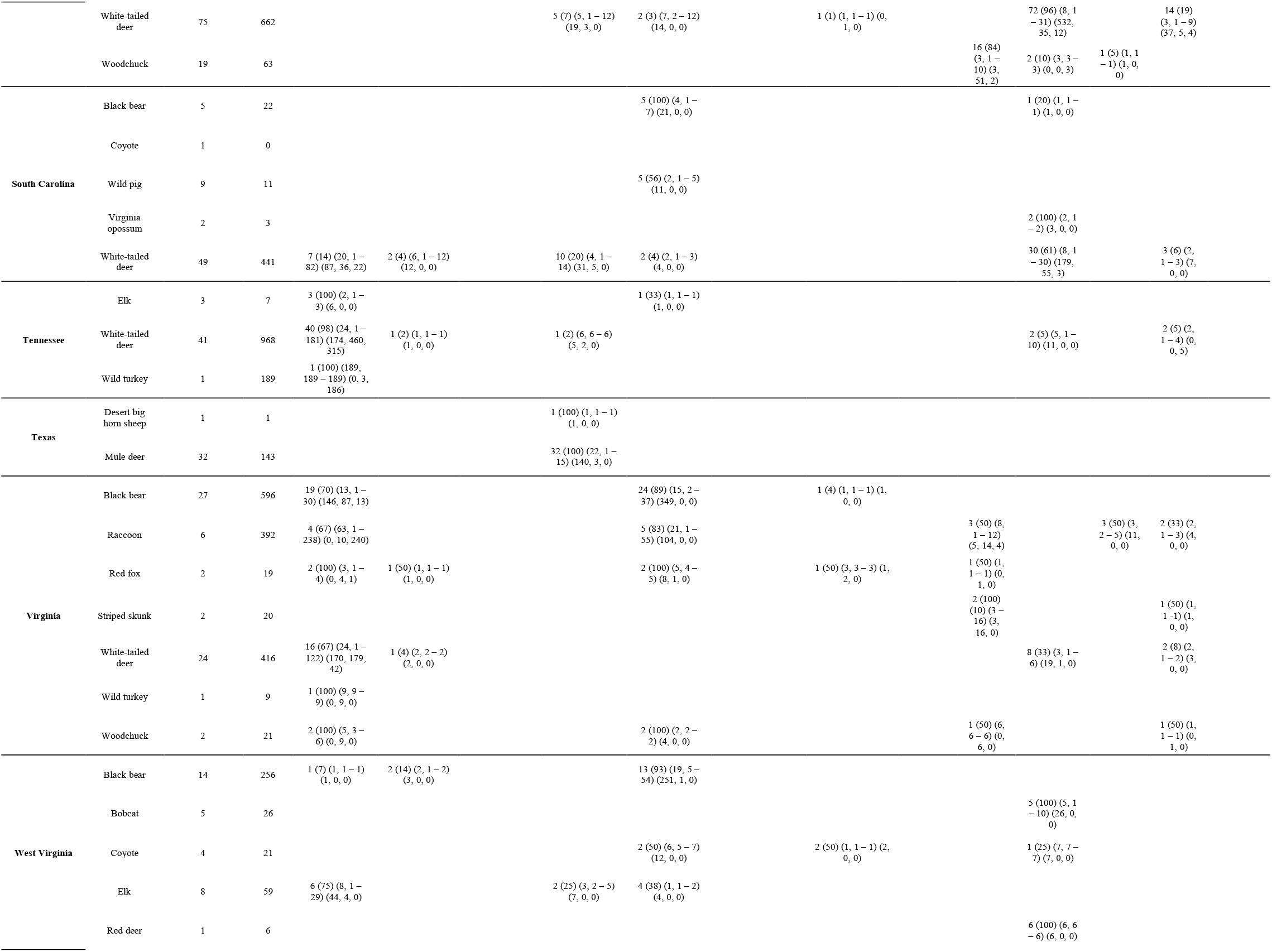
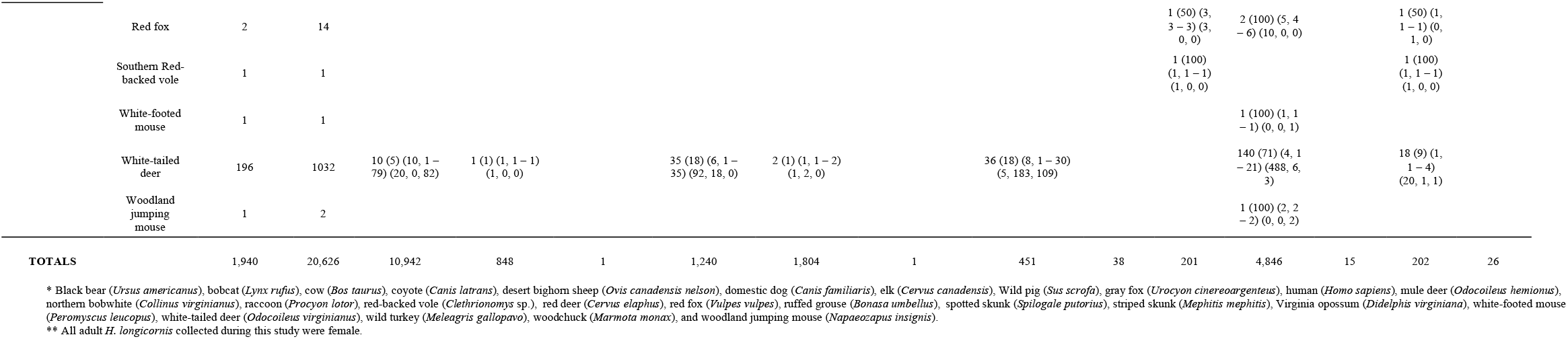

**SFigure 1.**
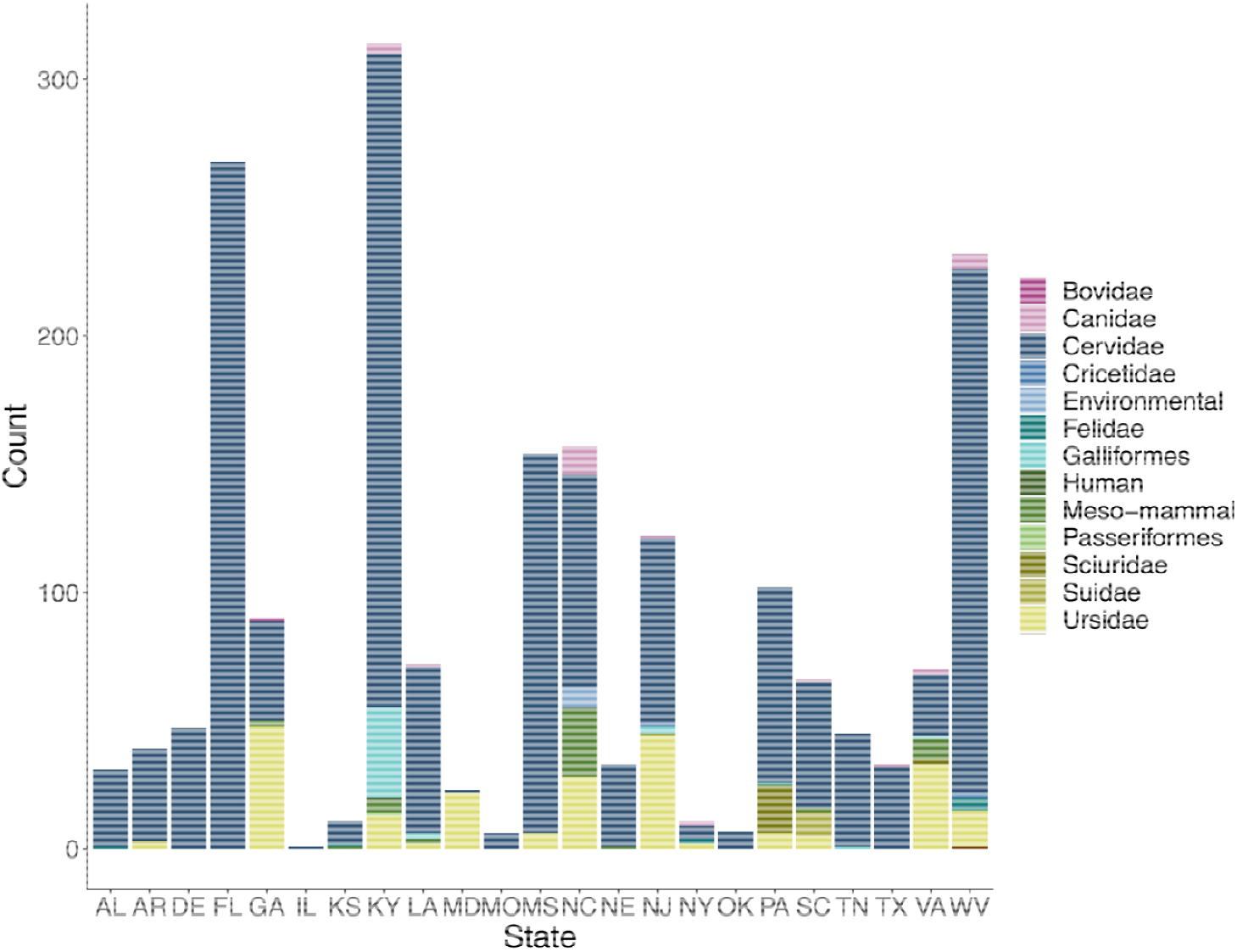
Host submissions by state, 2010 – 2021. Stacked bar chart represents total number of submissions by state, bars are colored in by number of host group submitted.

**SFigure 2.**
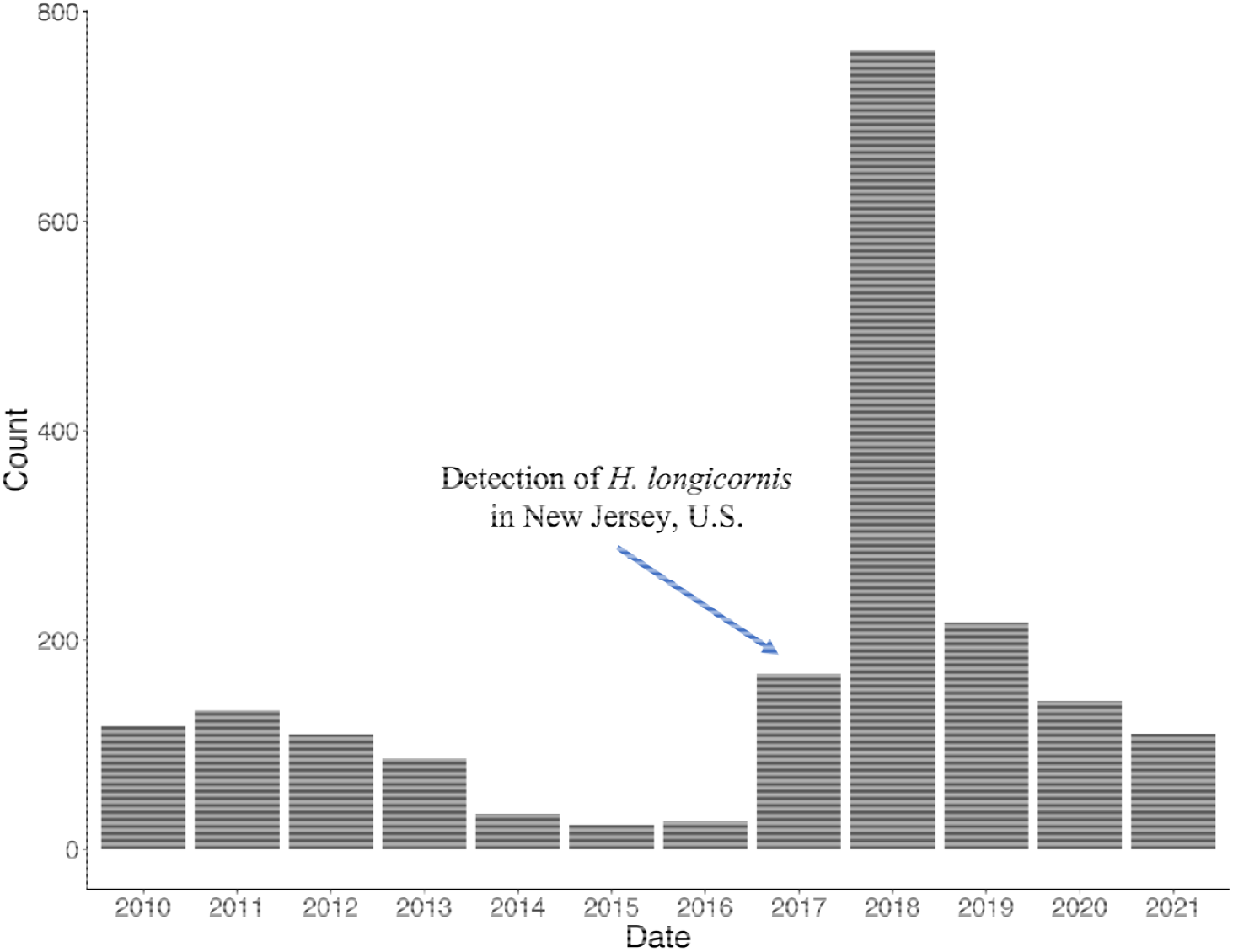
Number of submissions over the years during surveillance, 2010 - 2021.

